# Structural bases for the Charcot–Marie–Tooth disease induced by single amino acid substitutions of myelin protein zero

**DOI:** 10.1101/2023.04.12.536520

**Authors:** Masayoshi Sakakura, Mikio Tanabe, Masaki Mori, Hideo Takahashi, Kazuhiro Mio

**Affiliations:** Graduate School of Medical Life Science, Yokohama City University, Yokohama 230-0045, Japan; Structural Biology Research Center, Institute of Materials Structure Science, KEK/High Energy Accelerator Research Organization, Tsukuba 305-0801, Japan; AIST-UTokyo Advanced Operando-Measurement Technology Open Innovation Laboratory (OPERANDO-OIL), National Institute of Advanced Industrial Science and Technology (AIST), Kashiwa 277-0882, Japan

**Keywords:** Charcot–Marie–Tooth disease, compact myelin, myelin protein zero P0, membrane adhesion, homophilic protein-protein interaction

## Abstract

Myelin protein zero (MPZ or P0) is a major transmembrane protein expressed in peripheral compact myelin and functions to glue membranes to form multiple layered membranes characteristic of myelin. Intermembrane adhesion is mediated by homophilic interactions between the extracellular domains (ECDs) of MPZ molecules. Single amino acid substitutions in an ECD cause demyelinating neuropathy, known as Charcot–Marie–Tooth disease (CMT); however, the mechanisms by which such substitutions induce the disease are not well understood. To address this issue, we constructed a novel assay to evaluate the membrane-stacking activity of ECD using ECD-immobilized nanodiscs. Using this novel “nanomyelin” system, we found that octameric (8-meric) ECDs with a stacked-rings-like configuration are responsible for membrane adhesion. Two inter-ECD interactions, *cis* and head-to-head, are essential to constituting the 8-mer and, consequently, to gluing the membranes. This result was further reinforced by the observation that the CMT-related N87H substitution at the *cis* interface abolished membrane-adhesion activity. In contrast, the CMT-related D32G and E68V variants of ECD retained membrane-stacking activity, whereas their thermal stability was reduced compared to that of the WT. Reduced thermal stability may lead to impairment of the long-term stability of ECD and the layered membranes of myelin.

## Introduction

Myelin covering a neuronal axon functions as an electrical insulator that is essential for rapid signal conduction along the axon, known as saltatory conduction. In the peripheral nervous system (PNS), myelin is formed by Schwann cells that spiral around axons to form layered membranes. The membranes are regularly and densely packed in the region called compact myelin, where the spaces between extracellular membranes and between intracellular membranes are maintained to be approximately 50 and 30–40 Å, respectively (1). Myelin protein zero (MPZ or P0) is the most abundant protein expressed in the compact myelin of the PNS (2) and functions as an adhesion molecule to stack membranes (3). MPZ is essential for proper myelin formation, and genetic mutations leading to amino acid substitutions in MPZ cause inherited demyelinating neuropathies known as Charcot–Marie–Tooth disease (CMT) (4). MPZ is a type 1 membrane protein composed of an immunoglobulin-like extracellular domain (i.e., ECD), a single transmembrane helix, and an intracellular region. Reportedly, MPZ exists as a mixture of monomers, dimers, and tetramers; however, the biological significance of these multimers in myelin is poorly understood (5–7). Two crystal structures of ECD have been determined: an ECD structure from human MPZ (hMPZ-ECD) fused with maltose-binding protein (MBP) (8) and an ECD structure from rat MPZ (rMPZ-ECD) (6). HMPZ-ECD exists as a monomer in the crystal because of the presence of fused MBP, and the structure provides little information about the molecular interactions of ECD (8). In contrast, the rMPZ-ECD crystallized as a multimer, and three types of inter-ECD interactions were observed. The *cis* interaction formed an ECD tetramer by binding four ECDs with the same orientation. The *trans* and head-to-head interactions laterally and vertically connect the two ECDs in opposite directions, respectively (6). The *trans* and head-to-head interactions potentially adhere two membranes via ECD, and the former interaction has been proposed to be responsible for the compact myelin formation since the resultant ECD multimer fits the approximately 50 Å gap between the membranes histologically observed in myelin (6). Curiously, CMT-related amino acid substitution sites have been found at these three interaction sites (4), suggesting that all these sites are related to proper myelin formation. However, the significance of these substitutions in membrane adhesion has not yet been well examined, partially because cell biology or animal experiments are time-consuming and unsuitable for comparing the activities of many ECD variants (9, 10). Therefore, an alternative experimental tool for analyzing the activity of ECD variants must be developed.

In this study, we established a new method for analyzing the membrane-stacking activity of human MPZ-ECD (hMPZ-ECD; hereafter referred to as ECD) by monitoring the oligomerization of ECD-immobilized nanodiscs (NDs) (11); Using this “nanomyelin” system in combination with ECD variants with single amino acid substitutions, we identified the residues responsible for the membrane adhesion activity of ECD. We also found that some CMT-related amino acid substitutions did not affect the membrane-stacking activity of ECD but reduced the thermal stability of the protein. Based on these results, we provide molecular insights into CMT initiated by amino acid substitutions in the ECD.

## Results

### Oligomerization state of ECD in solution

We prepared a recombinant ECD with a C-terminal histidine-tag (ECD-His) by refolding and analyzed its multimeric states using size-exclusion chromatography (SEC) with a multi-angle light scattering (MALS) detector (Fig. 1). The protein eluted in two main peaks, and MALS revealed that it contained particles with apparent sizes of 126 (±2) kDa and 17 (±0.03) kDa, which correspond to octameric (8-meric) and monomeric ECD-His, respectively. We independently reapplied the second SEC analysis to the 8-meric and monomeric ECD-His eluates and confirmed that they produced two peaks, similar to the elution pattern of the first SEC (Fig. S1). These results indicated the existence of an 8-mer-monomer equilibrium for ECD-His. Notably, the retention volume of monomeric ECD-His (19–20 mL in Fig. 1) was larger than the expected volume of 16– 17 mL based on the molecular weight (MW) of ECD-His, indicating that monomeric ECD-His interacted with the gel matrix (agarose and dextran) of the SEC column.

**Figure 1.**
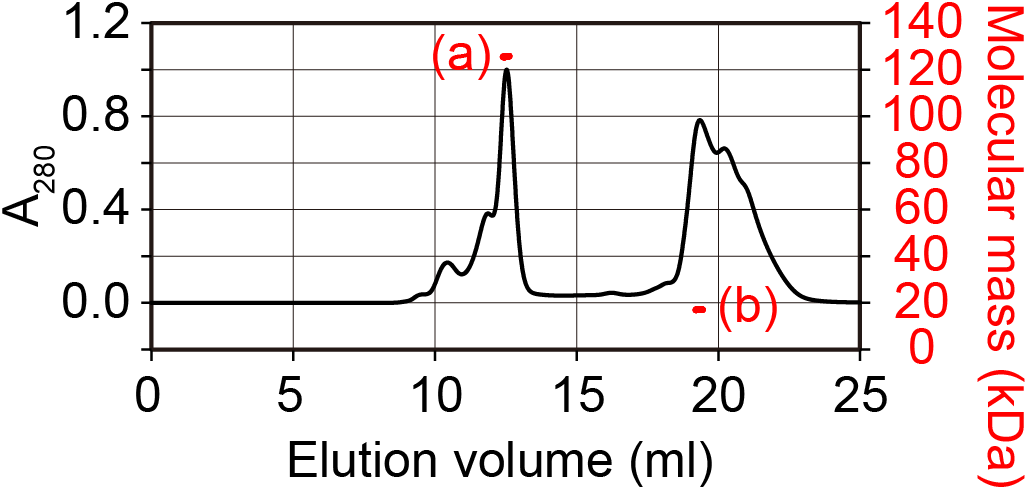
SEC-MALS profile of ECD-His. The black line indicates the SEC profile of ECD-His monitored by the absorbance at 280 nm, and the red lines show the molecular mass of the corresponding elution peaks. The molecular mass indicated by (a) and (b) correspond to 126 and 17 kDa, respectively. The concentration of the injected ECD-His was 52 μM. The experiment was performed in the standard buffer using a Superdex 200 Increase 10/300 column (Cytiva).

### Crystal structure of the multimeric human MPZ-ECD

We determined the crystal structure of ECD (without a histidine-tag) with a 2.1 Å resolution to obtain the molecular basis for the multimerization of human MPZ-ECD. The structure matched well with the previously reported structure of rMPZ-ECD with an RMSD value of 0.387 Å (115 C_α_ atoms in the amino acid residues 1–119 excluding the missing 4 C_α_s in rMPZ-ECD). The F-G loop (N102–G108), which was disordered in the rat structure, was visible in the human structure (Fig. S2) (6). The molecular contacts of human ECDs were similar to those of rat ECDs in the crystal, where each ECD contacted the other four ECDs through three types of inter-ECD interactions (Fig. 2A and S3). A ring-shaped ECD tetramer was formed through *cis* interactions, and the ring was laterally and vertically attached to other tetramers in the opposite direction via *trans* and head-to-head interactions, respectively (Fig. 2A). The residues involved in the inter-ECD interactions are shown in Fig. 2B–D. Aromatic residues play central roles in all the interfaces, as seen in the rMPZ-ECD structure: W24 makes a π-stacking interaction with H86 of the facing ECD at the *cis* interface (Fig. 2B), H52 forms a cation-π interaction with R45 at the *trans* interface (Fig. 2C), and W28 makes hydrophobic interactions with a cavity formed by D32 and K55 at the head-to-head interface (Fig. 2D). Detailed structural differences between human and rat MPZ-ECDs caused by the three amino acid substitutions are described in the Supplemental Results.

**Figure 2.**
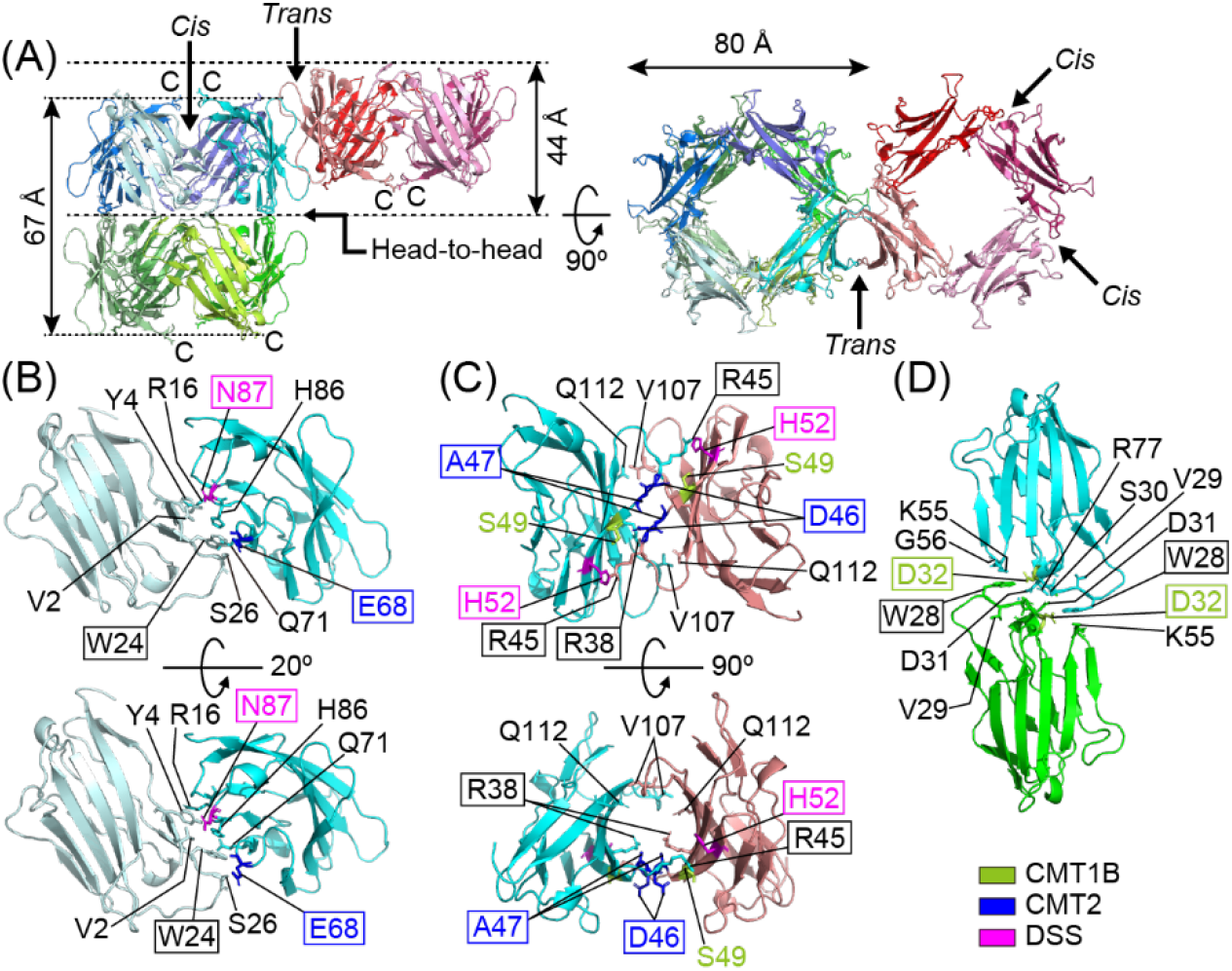
Crystal structure of multimerized human MPZ-ECDs. (A) Overview of the multimerized structure of ECDs. Three ring-shaped tetramers are shown in bluish, greenish, and reddish colors. In each of these rings, ECDs are connected by *cis* interactions. The *trans* interaction is formed between the laterally aligned ECDs in opposite directions, such as the cyan and salmon pink ECDs. The head-to-head interactions are formed between the vertically aligned ECDs, such as the cyan and green ECDs. The C-terminal ends of ECDs are shown by C. Putative positions of membrane surfaces are indicated by dotted lines. (B–D) Residues involved in the (B) *cis*, (C) *trans*, and (D) head-to-head interactions are labeled, and their side chains are shown with sticks. Residues related to CMT1B, CMT2, and DSS are colored lime green, blue, and magenta, respectively. Amino acid substitution sites examined in this study are boxed.

Two possible interactions are responsible for the membrane-stacking activity of the ECD. The *trans* interaction laterally connects two ECDs, whose C-terminus is linked to the transmembrane helix via a short linker in the native protein, and the *trans*-interacted ECDs separate the two membranes with a distance of more than 44 Å (inter-W28-C_ζ2_ distance along the membrane-plane normal vector) (Fig. 2A). The head-to-head interaction is the other option to glue the membranes, and the inter-membrane distance becomes more than 67 Å in this case (inter-E119-C_α_ distance along the membrane-plane normal) (Fig. 2A). The simultaneous formation of *trans* and head-to-head interactions is unlikely because it disturbs the planar structure of the membranes.

The oligomerized ECDs showed differently shaped B′-C (S25–I33) and F-G (N102–G108) loops from those of the monomeric ECD fused with MBP (8), although the overall ECD structures were similar with an RMSD of 1.550 Å (Fig. S4A and B). In the multimeric ECD, the B′-C loop is involved in both the *cis* and head-to-head interactions, and the F-G loop is involved in the *trans* interaction. In monomeric ECD, some residues on these loops form crystal contacts with the surrounding MBPs, so the resultant structure does not necessarily reflect the native monomeric conformation. Nevertheless, structural differences in these loops between the two ECD structures suggest that the B’-C and F-G loops are flexible, and structural changes occur upon the formation of inter-ECD interactions.

### In vitro assay to examine the membrane-stacking activity of ECD

To identify the residues that are responsible for the membrane-stacking activity of ECD, we developed an assay system to analyze the membrane-stacking activity of ECD by monitoring the oligomerization of ECD-immobilized nanodiscs (NDs) (11). We prepared NDs containing Ni^2+^-chelating lipids on which ECD-His molecules were bound (ECD-His-NDs) (Fig. S5). Fig. 3A shows the SEC profile of ECD-His-NDs. Fig. S6 shows the proteins present in each elution peak. ECD-His-NDs were eluted at the void volume (8.2 mL), which corresponded to a MW >1120 K. Because the MW of monomeric ECD(W24A)-His-ND was 300 K (described below; Fig. 4D), it was assumed that ECD-His-NDs would form at least a tetramer in the eluate. The multimerization of ECD-His-NDs was partially inhibited by the coexistence of ECDs (without a histidine-tag) and completely impaired by the addition of imidazole (Fig. 3B and S6), suggesting that multimerization is mediated by interactions between ECD-His attached to the lipids.

**Figure 3.**
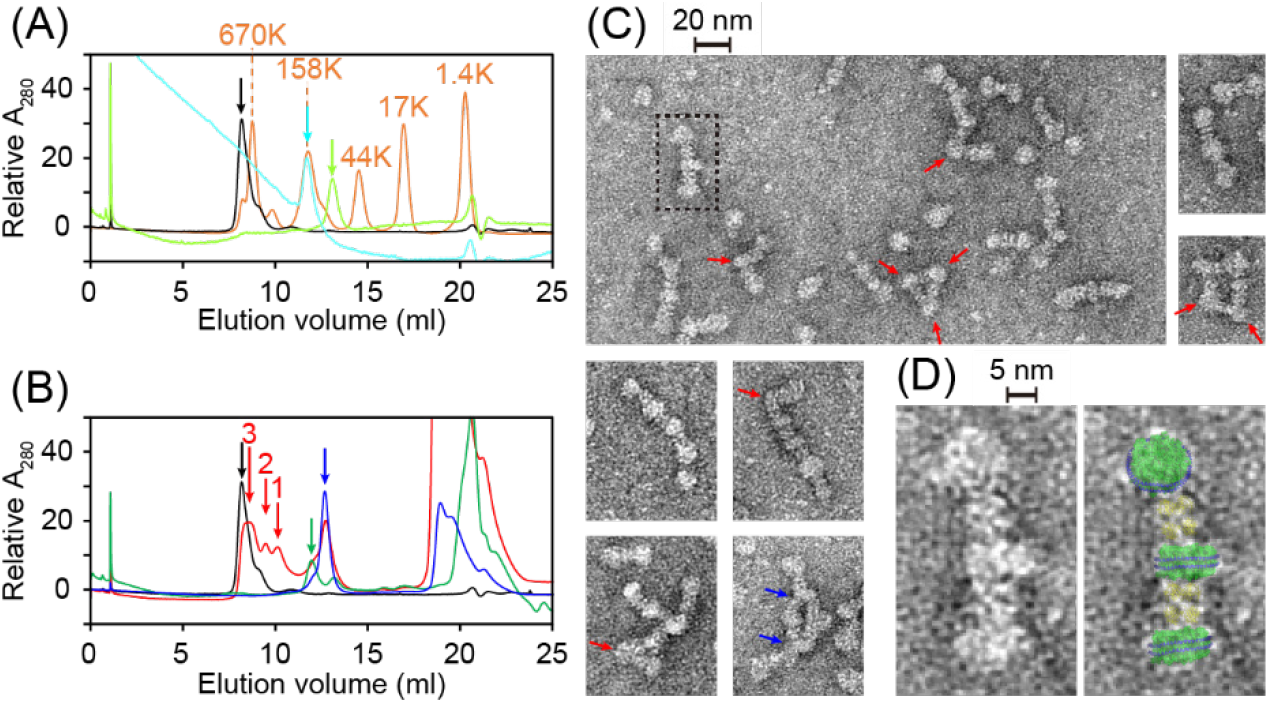
Membrane-stacking activity of ECD-His. (A) SEC profiles of ECD-His (lime green), ND (cyan), their mixture (black), and MW standards (orange). Lime green and cyan arrows indicate the elution peaks of the 8-meric ECD-His and monomeric ND, respectively. The ECD-His-ND complex eluted at the void volume (black arrow), indicating more than four NDs are connected by ECD-His. (B) Inhibition experiments to identify the interactions essential for the ND-multimerization. The SEC profiles of the ECD-His-ND (black), the mixture of the ECD-His-ND and an excess amount of ECD without a histidine-tag (red), and the mixture of the ECD-His-ND and 250 mM imidazole (green) are shown. The blue line shows the profile of ECD without a histidine-tag. The peak originating from the multimerized ECD-His-ND (black arrow) was attenuated or disappeared by the addition of ECD or imidazole, respectively, indicating that NDs were linked by the homophilic interactions between ECD-His and by the interaction between ECD-His and Ni^2+^-chelating lipids in NDs. Peaks with red arrows correspond to the MWs of the monomeric (arrow 1), dimerized (arrow 2), and trimerized (arrow 3) ECD-His-NDs, respectively. Green and blue arrows indicate the elution peaks of the monomeric ND and 8-meric ECD without a histidine-tag, respectively. The SEC analyses were performed using a Superdex 200 Increase 10/300 column (Cytiva) in the standard buffer. (C) Representative EM images of the ECD-His-NDs. The SEC eluate at the void volume was used for the observation. Large particles with the size equivalent to NDs are connected linearly by small particles with the size of the stacked-rings-ECD-8-mer. Turns and branches are shown by red and blue arrows, respectively. (D) Expanded view of a representative rod-like structure in the dotted box in (C) without (left) and with (right) manually superimposed structural images of the 8-meric ECDs and NDs. Structural coordinates of an MSP1D1-utilizing ND were taken from PDB (ID 7RSC). The ECDs and NDs are colored as Fig. S5.

**Figure 4.**
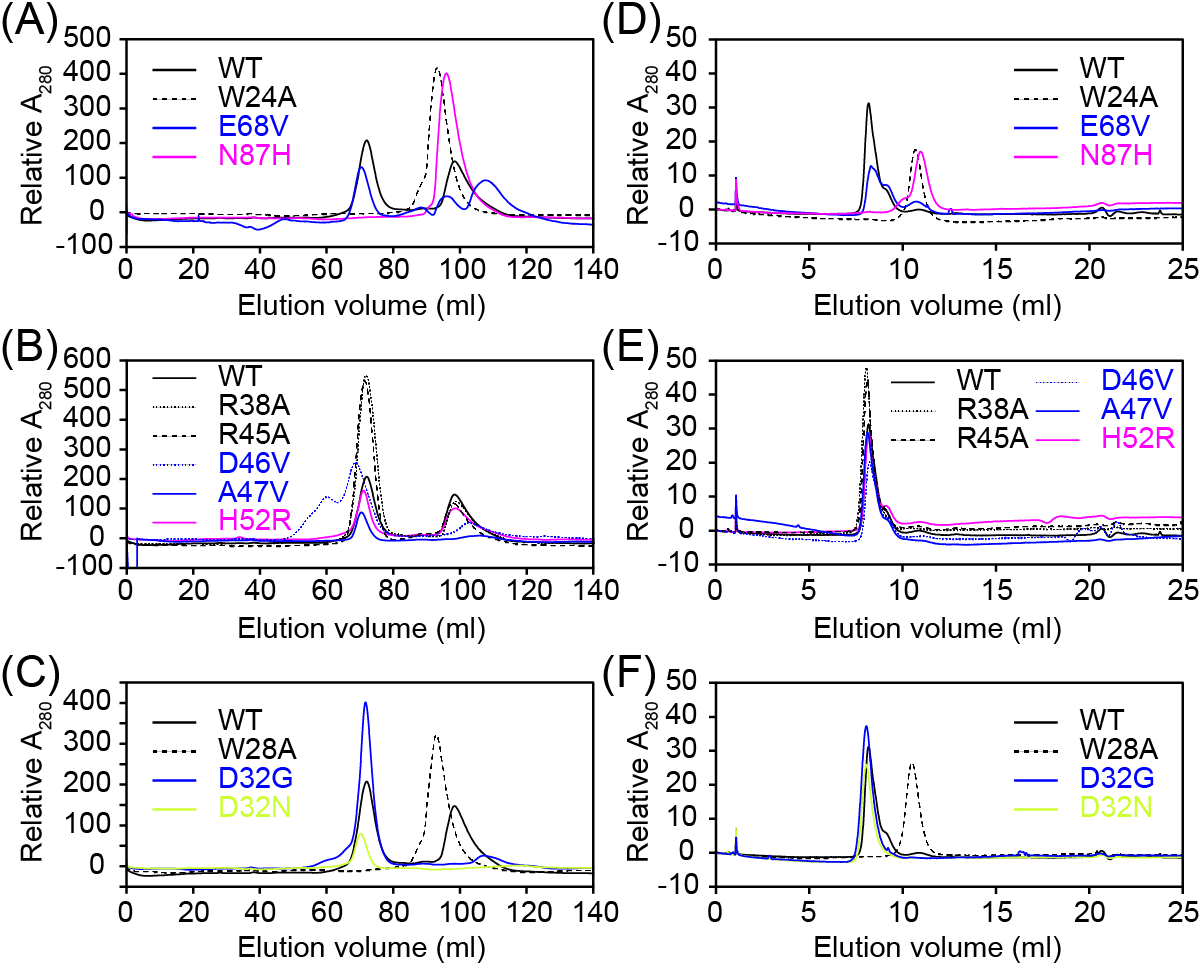
Multimerization and membrane-adhesion activities of ECD-His-variants. SEC profiles of the ECD-His variants with substitutions at the (A) *cis* (W24A, E68V, and N87H), (B) *trans* (R38A, R45A, D46V, A47V, and H52R), and (C) head-to-head (W28A, D32G, and D32N) interfaces. The analyses were performed in the standard buffer using a HiLoad Superdex 200 16/600 column (Cytiva). Most variants eluted as a mixture of an 8-mer and the monomer, while W24A, N87H, and W28A eluted solely as the monomer. SEC profiles of the mixtures of ND and the ECD-His variants with substitutions at the (D) *cis*, (E) *trans*, and (F) head-to-head interfaces. The analyses were performed in the standard buffer using a Superdex 200 Increase 10/300 column (Cytiva). The W24A and N87H substitutions at the *cis* interface and the W28A substitution at the head-to-head interface completely impaired the multimerization of NDs. None of the 5 substitutions at the *trans* interface affected the multimerization. Results of the variants with CMT1B-, CMT2-, and DSS-related substitutions are colored lime green, blue, and magenta, respectively.

Fig. 3C shows electron micrographic (EM) images of the ECD-His-NDs contained in the SEC eluate. Rod-like structures with alternating thick and thin parts were observed at regular intervals in each rod, and turns or branches were occasionally observed (arrows in Fig. 3C). The average height/width of the thicker parts was 71 (±11)/108 (±12) Å, and that of the thinner parts was 83 (±9)/74 (±9) Å. Fig. 3D shows an expanded view of an EM image of a typical rod-like structure where structural images of an ND (height/width = 50/97 Å (11)) and the 8-meric ECDs assembled by the *cis* and head-to-head interactions (height/width = 67/80 Å, Fig. 2A) were manually superimposed to guide the size of the particles. The sizes of the thicker and thinner parts roughly correspond to those of the ND and 8-meric ECDs, respectively, suggesting repetitive assembly of the NDs and 8-meric ECDs (Fig. S5). Because these stacked NDs connected by ECD 8-mers mimic the layered membranes glued by MPZ in myelin, hereafter, we describe them as nanomyelin. The turns and branches of nanomyelin may be caused by a tilted ECD-8-mer attached to the edge of an ND via one or two histidine-tags and by the attachment of multiple ECD-8-mers on the surface of a single ND, respectively.

### The *cis* and head-to-head interactions were responsible for the membrane-stacking

To identify the inter-ECD interaction site(s) responsible for nanomyelin formation, we prepared four representative ECD-His variants with alanine substitutions at the residues involved in inter-ECD interactions in the crystal structure (W24A at the *cis* interface, R38A and R45A at the *trans* interface, and W28A at the head-to-head interface) (Fig. 2B–D).

Using these ECD variants, we analyzed their multimerization by SEC and their membrane-stacking activity using nanomyelin experiments. Fig. 4 shows the SEC profiles of the ECD-His variants (Fig. 4A–C) and ECD-His-variant-immobilized NDs (ECD-His-variant-NDs, Fig. 4D–F). Fig. 5 shows the EM images of the main peak fraction from the SEC of the ECD-His-variant-NDs. SEC experiments revealed that the W24A (a *cis* interface substitution) and W28A (a head-to-head interface substitution) variants existed solely as monomers (Fig. 4A and C) and that these variants did not induce multimerization of NDs (Fig. 4D, 4F, 5A, and 5I). In contrast, the two variants with substitutions at the *trans* interface (R38A and R45A) behaved as the WT; each existed as a mixture of an 8-mer and the monomer (Fig. 4B) and induced ND-multimerization to form nanomyelin (Fig. 4E, 5D, and 5E). These results indicate that (1) W24 and W28 play critical roles in the *cis* and head-to-head interactions, respectively; (2) ECDs form an 8-mer assembled by the *cis* and head-to-head interactions in solution; (3) both *cis* and head-to-head interactions are required for the formation of the 8-mer and the membrane-stacking activity of ECD; and (4) the *trans* interaction is unlikely to be formed in solution and does not contribute to nanomyelin formation.

**Figure 5.**
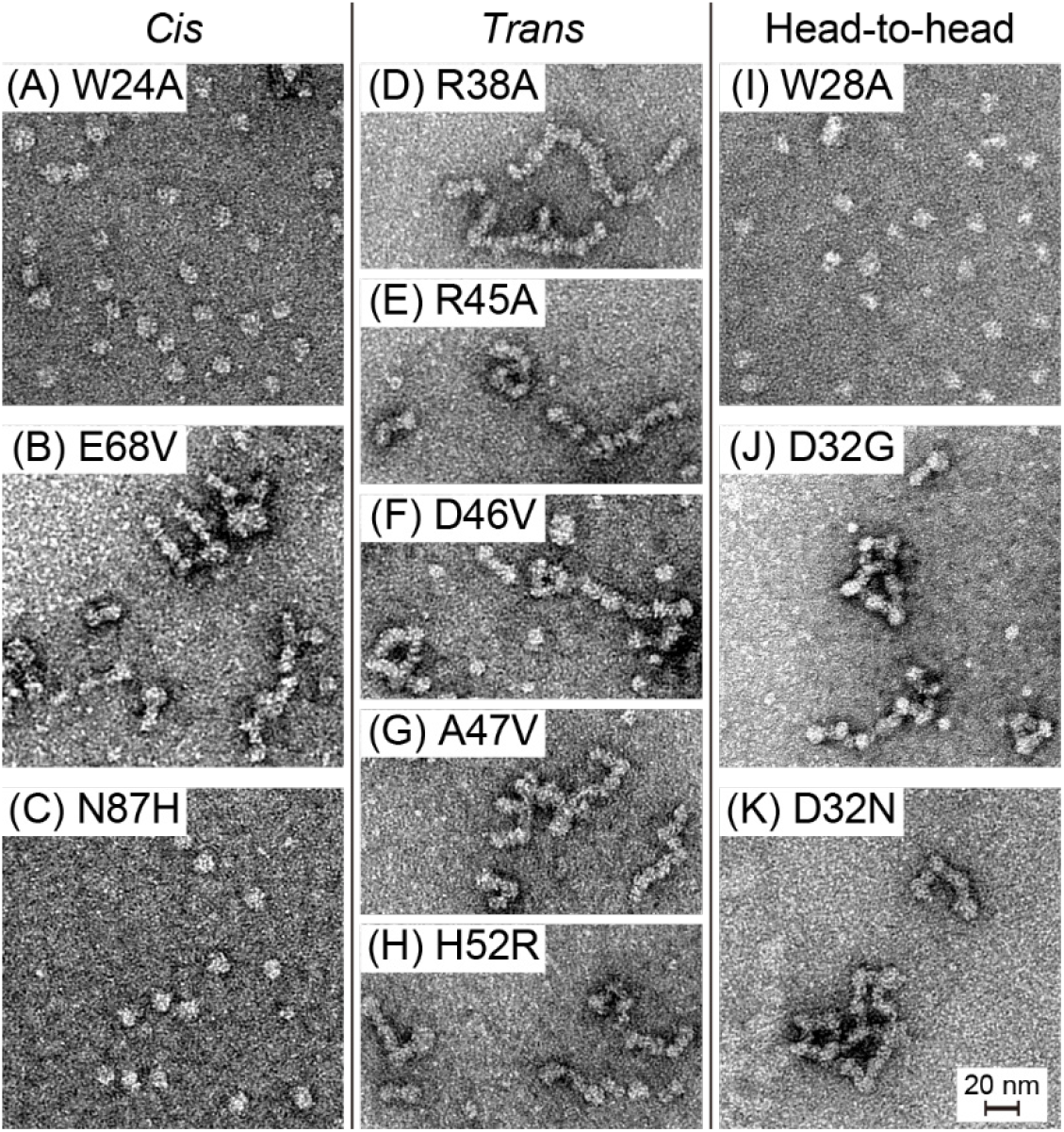
Effects of the amino acid substitutions of ECD-His on the ND-assembly. EM images of the NDs complexed with the ECD-His variants with substitutions at the *cis* interface (A, B, and C correspond to W24A, E68V, and N87H, respectively), *trans* interface (D, E, F, G, and H correspond to R38A, R45A, D46V, A47V, and H52R, respectively), and head-to-head interface (I, J, and K correspond to W28A, D32G, and D32N, respectively) are shown. The W24A, N87H, and W28A variants did not assemble NDs, indicating these residues are important to stack NDs. None of the 5 substitutions at the *trans* interface affected the ND-assembly. Figure 6. Thermal stability of ECD variants. Thermal denaturation curves of the ECD-His variants with amino acid substitutions at the (A) *cis*, (B) *trans*, and (C) head-to-head interfaces. Fractional changes in the molar ellipticity at 217 nm are plotted against the temperature for each variant. Lime green, blue, and magenta indicate the results of the CMT1B-, CMT2-, and DSS-related ECD-variants, respectively.

**Figure 6.**
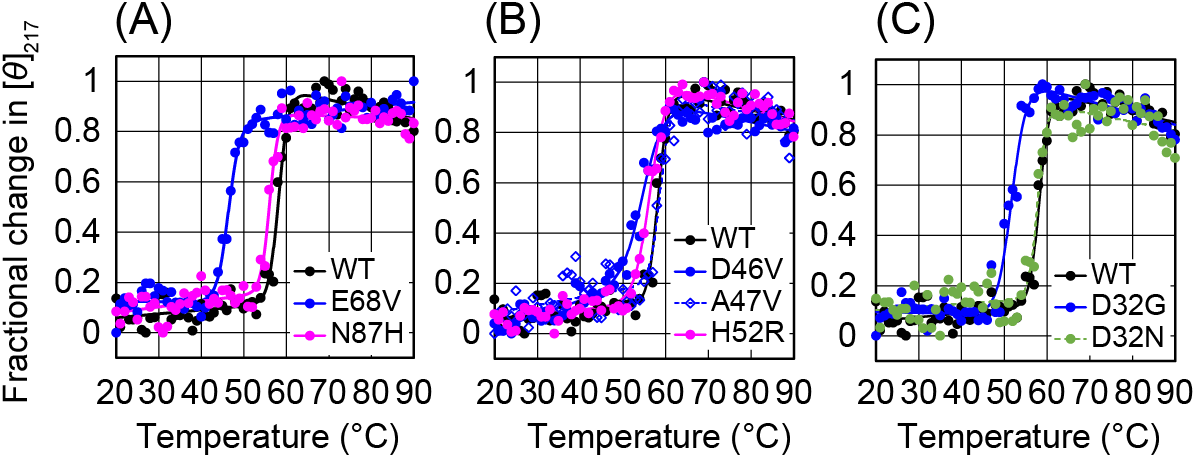
Thermal stability of ECD variants. Thermal denaturation curves of the ECD-His variants with amino acid substitutions at the (A) *cis*, (B) *trans*, and (C) head-to-head interfaces. Fractional changes in the molar ellipticity at 217 nm are plotted against the temperature for each variant. Lime green, blue, and magenta indicate the results of the CMT1B-, CMT2-, and DSS-related ECD-variants, respectively.

### Effects of the CMT-related amino acid substitutions on the membrane-stacking activity of ECD

We prepared ECD-His variants with CMT-related substitutions (E68V and N87H at the *cis* interface; D46V, A47V, and H52R at the *trans* interface; and D32G and D32N at the head-to-head interface) to determine the effects of these substitutions on nanomyelin formation (Fig. 2B–D).

Among disease-related substitutions, H52R and N87H are linked to Dejerine–Sottas Syndrome (DSS; a severe type of demyelinating neuropathy), and D32G, D46V, A47V, and E68V are causative of CMT Type 2 (CMT2) (4). The D32N substitution produces an additional glycosylation site on ECD, leading to CMT Type 1B (CMT1B) (12). The recombinant protein used in this study was non-glycosylated. Therefore, the results obtained for the D32N variant solely reflected the effect of side-chain neutralization.

As shown in Fig. 4 and 5, the N87H variant with a *cis* interface substitution existed as a monomer (Fig. 4A) and did not induce the multimerization of NDs (Fig. 4D and 5C). In contrast, E68V (a *cis* interface substitution), D32G and D32N (head-to-head interface substitutions), and three variants with substitutions at the *trans* interface (D46V, A47V, and H52R) existed as a mixture of an 8-mer and a monomer (Fig. 4A–C) and induced ND multimerization to form nanomyelin (Fig. 4D–F, (5B, 5F–H, 5J, and (5K). These results indicate that (1) the N87H substitution impairs the membrane-stacking activity of ECD by destroying the *cis* interaction and (2) the D32G, D32N, D46V, A47V, H52R, and E68V substitutions do not affect the inter-ECD interactions or membrane adhesion activity of ECD.

### Thermal stability of the ECD variants

In the nanomyelin experiments, the amino acid substitutions D32G, D32N, D46V, A47V, H52R, and E68V did not impair the membrane-stacking activity of ECD, although these substitutions induced CMT. To examine the effect of these substitutions on the thermal stability of ECD, we analyzed the temperature-dependent denaturation of the ECD-His variants using circular dichroism (CD) spectroscopy (Fig. 6). For all ECD-His variants, the protein samples precipitated in the heating processes, and the reversed cooling processes did not show apparent changes in [*θ*]_217_. These results indicate that the observed processes were irreversible. The data were fitted to a theoretical curve based on a two-state fold-unfold equilibrium (13) to obtain the midpoint temperatures (T_m_s) for denaturation. The T_m_ values are listed in Table 1. The E68V (*cis* interface; Fig. 6A) and D32G (head-to-head interface; Fig. 6C) substitutions reduced T_m_ by 11 and 6 °C from the WT value of 58 °C, respectively. On the other hand, the N87H (*cis* interface), D32N (head-to-head interface), and three variants with substitutions at the *trans* interface (D46V, A47V, and H52R) showed marginal (< 3 °C) differences in T_m_ from WT (Fig. 6).

**Table 1.**
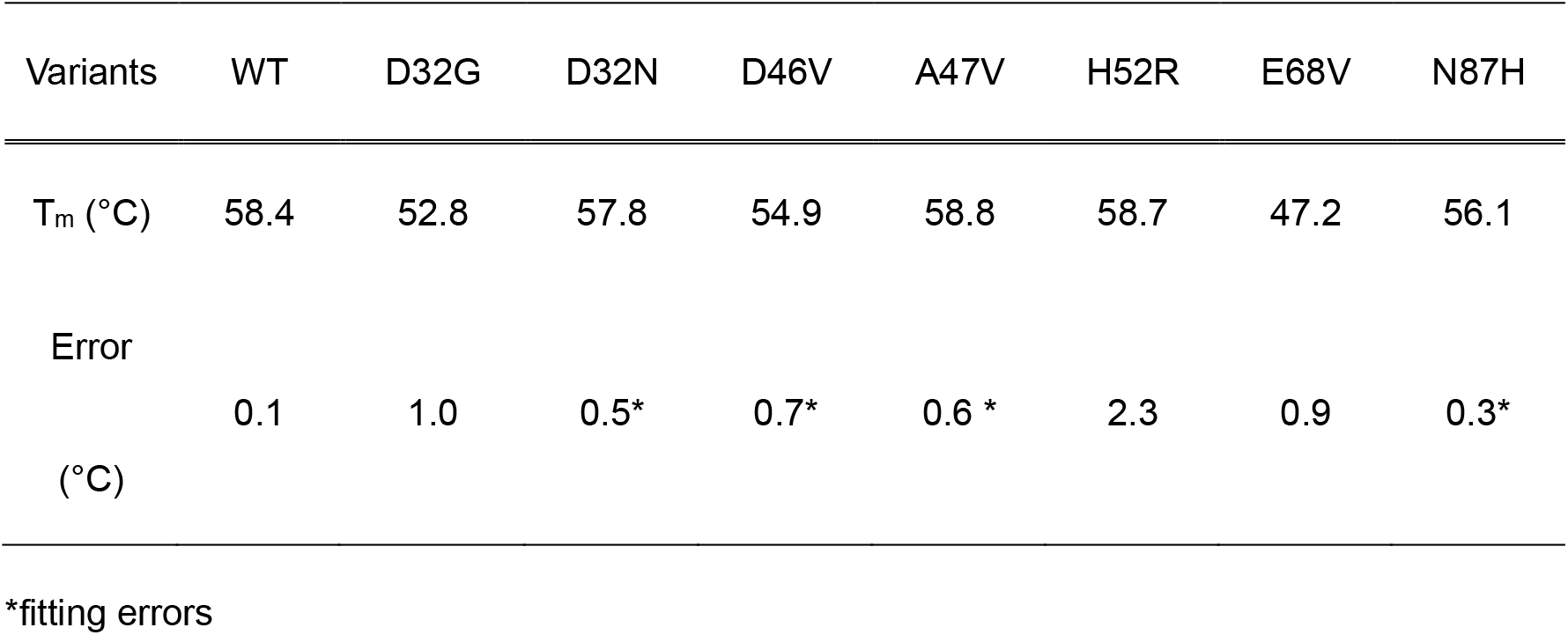
Midpoint temperatures of the denaturation (T_m_s) of the ECD-His variants

| Variants | WT | D32G | D32N | D46V | A47V | H52R | E68V | N87H |
| --- | --- | --- | --- | --- | --- | --- | --- | --- |
| $T_m$ (°C) | 58.4 | 52.8 | 57.8 | 54.9 | 58.8 | 58.7 | 47.2 | 56.1 |
| Error<br>(°C) | 0.1 | 1.0 | 0.5* | 0.7* | 0.6 * | 2.3 | 0.9 | 0.3* |

\*fitting errors

## Discussion

### Inter-ECD interactions responsible for the membrane-stacking

Based on the previously reported crystal structure of multimeric rMPZ-ECD (6), the *trans* interaction, which laterally combines ECDs in opposite directions, has been considered responsible for the formation of membrane layers in compact myelin (14). In this study, we confirmed that the *trans* interaction was formed in the crystal (Fig. 2), but this *trans* interaction was unlikely to be formed in solution because the variants with substitutions at the *trans* interaction site did not affect 8-mer formation or nanomyelin formation (Fig. 4 and 5). Rather, ECDs in solution formed *cis* and head-to-head interactions to construct stacked-rings like 8-mer (the bluish and greenish ECDs in Fig. 2A, hereafter the stacked-rings-8-mer), and this 8-mer was responsible for membrane-stacking because some amino acid substitutions at these interfaces impaired the oligomerization and membrane-adhesion activities of ECD (Fig. 3–5). An emerging issue here is that the 67 Å vertical length of the stacked-rings-8-mer exceeds the known extracellular inter-membrane distance of 47 Å in compact myelin (15), and the 8-mer does not seem to fit the inter-membrane gap. A possible explanation for this is that the stacked-rings-8-mer plays a role in the early stages of myelination. It has been reported that myelin-associated glycoproteins (MAGs) primarily attach the mesaxon membranes in the myelinating Schwann cells with the inter-membrane distance of approximately 120 Å, and these loosely layered membranes are eventually compacted to make the inter-membrane distance of 47 Å by replacing the adhesion proteins from MAG to MPZ (16). The stacked-rings-8-mer with the height of 67 Å may be formed during the myelin compaction to facilitate the transition of the inter-membrane gap from the MAG-mediated 120 Å to the final 47 Å which, in this case, would be achieved by the *trans* interaction between ECDs.

In contrast, a recent X-ray diffraction study reported that the extracellular inter-membrane distance in compact myelin is not uniform and varies from 40 to 65 Å (17). Taking into account the size of the polar heads of lipids (18), the stacked-rings-8-mer may be accommodated in the compact myelin area with a relatively large intermembrane distance and work to stabilize the membrane layers. This is consistent with the freeze-fractured image of compact myelin, in which particles with an averaged diameter of 86 Å, of which 50–70% would be MPZ (2), are distributed randomly on the membrane surface, with no lattice-like regular distribution of MPZ required for the formation of the *trans* interaction (19, 20).

In summary, we do not exclude the possibility that the *trans* interaction between ECDs stabilizes the membrane layers in compact myelin; however, we propose that the stacked-rings-8-mer of ECDs would also contribute to glue membranes, either as transient membrane stackers during the compaction process of PNS myelin or as major adhesion molecules in compact myelin regions with relatively longer intermembrane distances.

### Mechanism by which the N87H substitution extinguishes the membrane-stacking activity of ECD

The N87H substitution was found in a patient with Dejerine–Sottas syndrome (DSS), a severe form of demyelinating neuropathy, with three simultaneous amino acid substitutions in ECD: I85T, N87H, and D99N (21). In this study, we found that the N87H substitution induces monomerization of ECD and impairs its membrane-stacking activity (Fig. 4A, 4D, and 5C). In the crystal structure of the multimeric ECDs, N87 was located at the *cis* interface and faced V2 of the adjacent ECD (Fig. 2B). The distance between the C_β_ of N87 and the C_γ2_ of V2 is 4.2 Å, indicating these atoms form a van der Waals interaction. The introduction of a bulk imidazole ring at N87 would induce a steric crash with V2 on the facing ECD, preventing the formation of *cis* interactions. In addition to the *cis* interaction, the N87H substitution prevents head-to-head interactions and causes monomerization of the protein. The W24A substitution at the *cis* interface also induced monomerization (Fig. 4A), suggesting that *cis* and head-to-head interactions are linked. In the crystal structure of the multimeric ECD, the B′-C loop constitutes both the *cis* and head-to-head interaction sites, and the structure of the loop is different from that of the monomeric ECD (Fig. S4A) (8). The *cis* interaction-induced conformational change of the B′-C loop would be a requisite for the formation of the head-to-head interaction. The head-to-head interaction in turn will stabilize the conformation of the B′-C loop to strengthen the *cis* interaction because the W28A substitution at the head-to-head interface also induces the monomerization of ECD (Fig. 4C). A subsequent kinetic study is required to fully understand the multimerization process of ECD. Notably, the transmembrane helix of MPZ contains a glycine-zipper oligomerization motif, which may stabilize the dimer and tetramer of intact MPZ, although we did not observe dimeric or tetrameric forms of ECD in the SEC experiment (22).

The N87H substitution induced a major loss-of-function effect on ECD, whereas DSS caused by heterozygous *MPZ* mutations have been associated with dominant-negative cytotoxic effects induced by aberrantly folded variant proteins (21, 23). Since the in vitro folding efficiency of the N87H variant observed in this study was higher than that of WT ECD, the N87H substitution may have a minor impact on the folding property of MPZ and work synergistically with the other substitutions (I85T and D99N) to cause the severe phenotype of this patient. To understand the role of the N87H substitution in DSS, an integrated analysis with the other substitutions may be necessary.

### Mechanism by which the D32G and E68V substitutions reduce the thermal stability of ECD

The D32G and E68V variants did not show apparent changes in membrane-stacking activity (Fig. 4 and 5); however, they exhibited reduced thermal stability relative to WT ECD (Fig. 6 and Table 1). These changes in thermal stability would result from changes in the unfolding process of the ECD-monomer, and the 8-mer-monomer transition will not be reflected in the changes in [*θ*]_217_ because changes in the β-sheet structure upon the 8-mer formation are subtle (β strand-fractions are 47 and 50% for the multimer and monomer, respectively).

E68 was located at the *cis* interface and faced S26 of the interacting ECD (Fig. 2B and S4A). These residues are 3.7 Å apart (between E68-C_α_ and S26-C_β_) and make a van der Waals interaction, while no polar interaction is formed within these residues. The E68V substitution retains the van der Waals contact with S26, so it is convincing that the substitution did not alter ECD oligomerization (Fig. 4A). Rather, the E68V substitution exposed the hydrophobic side chain on top of the C″-D loop to the solvent (Fig. S2), which may facilitate partial unfolding of the loop and subsequent denaturation of the entire protein.

D32 is located at the head-to-head interface, and its O_δ1_ is at a 3.6 Å distance from the N_ζ_ of K55 (Fig. 2D and S4C). The side-chains of these two residues are connected by an electrostatic interaction and form the wall of the cavity that accommodates the indole ring of W28 of the interacting ECD (Fig. S4C). D32G seems to destabilize this cavity and prevent the 8-merization of ECD by erasing the electrostatic interaction with K55; however, the variant formed an 8-mer and retained membrane adhesion activity, suggesting that the electrostatic interaction between D32 and K55 makes little contribution to the head-to-head interaction (Fig. 4C, 4F, and 5J). Rather, the electrostatic interaction may be important for stabilizing the β-sheet structure by bridging the edges of the strands C and C′ (Fig. S4C), and loss of the interaction by the D32G substitution may facilitate transient partial unfolding (the breathing motion) of β-sheet (24), which would contribute to lowering the T_m_. The fact that the D32N substitution did not affect the T_m_ may indicate that the hydrogen bond formed between N32 and K55 compensated for the electrostatic interaction.

## Conclusive remarks

We showed that hMPZ-ECDs form stacked-rings-8-mer via *cis* and head-to-head interactions in solution and that this 8-mer is responsible for the membrane-stacking activity of ECD. It is unclear whether the stacked-rings-8-mer with a 67 Å height forms transiently during myelin compaction or stably in compact myelin regions with an average inter-membrane extracellular gap of 47 Å. Further studies using Schwann cells or mutant animals are required to clarify this issue.

The N87H substitution, which is one of three simultaneous amino acid substitutions leading to DSS, induced pronounced effects on ECD activity; it eliminated multimerization and membrane-stacking activities. N87 is located at the *cis* interface and is not directly related to membrane adhesion. However, the N87H substitution allosterically affected the head-to-head interaction through the conformational change of the B′-C loop, which led to the loss of the membrane-stacking activity (Fig. S4A). This suggests that chemical compounds that bind to the *cis* interface may allosterically affect the head-to-head interface and revive membrane-stacking activity.

The effects of the CMT2-related D32G and E68V substitutions at the head-to-head and *cis* interfaces, respectively, were relatively moderate; they retained 8-mer-forming and membrane-stacking activities, while their thermal stabilities were lower than that of the WT. The reduced thermal stability of these variants may influence the long-term stability of these proteins in myelin and trigger the onset of CMT at middle-to-late ages (25, 26).

In this study, amino acid substitutions at the *trans* interface, including three disease-related substitutions (D46V, A47V, and H52R), did not result in any apparent changes in ECD activity. Our results do not necessarily exclude the possibility that this interaction occurs in compact myelin to stack membranes; however, another possibility is that ECD interacts with other myelin proteins essential for myelin compaction via this surface.

## Materials and methods

### Expression and purification of recombinant ECDs

The synthesized gene encoding human MPZ-ECD (I1 through V121), optimized for *Escherichia coli* expression, was obtained from ATUM and inserted into the pET-28a vector (Merck) using the restriction sites for Nco1 and Xho1. The resulting vector, pET-28a-hMPZ-ECD, produced ECD with two artificial amino acids (MG) at its N-terminus. Using the protocol attached to the QuikChange Site-Directed Mutagenesis Kit (Agilent), the vector pET-28a-hMPZ-ECD was modified to pET-28a-hMPZ-ECD-His, in which a hexa-histidine-tag (His_6_-tag) was linked to the C-terminus of the ECD via a linker with the PTR sequence. The amino acid sequences of these proteins are listed in Table S1. Vectors expressing variant proteins with single amino acid substitutions (W24A, W28A, D32G, D32N, R38A, R45A, D46V, A47V, H52R, E68V, and N87H) were produced using the protocol attached to the QuikChange Site-Directed Mutagenesis Kit (Agilent). WT and variant proteins were prepared using the protocol described below, unless otherwise noted.

Transformants were selected on LB plates with 30 μg/mL kanamycin after the plasmids were introduced into *E*. *coli* BL21(DE3). Two colonies were picked, and each colony was inoculated into 5 or 10 mL of LB medium with 30 μg/mL kanamycin in a 50 ml plastic tube (pre-culture). The pre-cultures were shaken at 37 °C overnight, and all of them were added to 0.25 L or 1 L of LB medium with 30 μg/mL kanamycin in a 1 L or 2 L baffled flask, respectively (main culture). The main culture was shaken at 37 °C and 190–210 rpm till the OD_600_ reached 0.7–1.3 for 2–3 h, and IPTG was added to a final concentration of 1 mM. The culture was kept shaking at 37 °C and 190–210 rpm overnight until the OD_600_ reached 4–6. The culture was centrifuged at 9,086 ×g and 4 °C for 20 min to spin down the cells. The cells were resuspended in a lysis buffer containing 25 mM Tris-HCl (pH 8.0), 150 mM NaCl, and 0.3% w/v CHAPS at a ratio of 20 mL of the lysis buffer to 1 g of wet cells. The cells were frozen in liquid nitrogen and stored at −80 °C until use. The wet cell weights were 3–8 g per 1 L culture.

The cells were lysed by sonication with repetitions of a 5 sec application and a 5 sec interval for 20 min. The resultant cell lysate was centrifuged at 27,143 ×g and 4 °C for 20 min. The ECD and ECD-His proteins were included in the precipitation. In the case of ECD (without a histidine-tag), the above precipitation was resuspended in Tris buffer 1 containing 25 mM Tris-HCl (pH 8.0), 1 mM EDTA, and 1% v/v Triton X-100 (in the ratio of 25 mL of the buffer to 1 g wet cells), sonicated for 5 min, and centrifuged at 27,143 ×g for 20 min at 4 °C. This wash procedure was repeated three times, and in the second and third washes, Tris buffer 2 containing 25 mM Tris-HCl (pH 8.0), 1 mM EDTA, and 1 M NaCl, and Tris buffer 3 containing 25 mM Tris-HCl (pH 8.0) and 1 mM EDTA, were used, respectively. The last suspension in Tris buffer 3 was centrifuged in plastic tubes at 7,600 ×g for 30 min at 4 °C. The ECD(-His) precipitate could be stored at −80 °C until use.

The ECD proteins were resuspended in solubilization buffer 1, which contains 8 M urea, 40 mM Tris-HCl (pH 8.2), 100 mM NaCl, and 1 mM 2-mercaptoethanol, at a ratio of 12.5 mL of the buffer to the washed inclusion body from 1 g of wet cells, and were rotated at room temperature overnight to solubilize the protein. ECD-His proteins were resuspended in solubilization buffer 2 containing 8 M urea, 40 mM Tris-HCl (pH 8.2), 100 mM NaCl, 1 mM EDTA, and 1 mM 2-mercaptoethanol at a ratio of 25 mL of the buffer to the inclusion body from 1 g of wet cells and rotated at room temperature overnight to solubilize the protein. The solution was centrifuged at 27,143 ×g and 25 °C for 30 min. ECD-His was purified using Ni^2+^ affinity chromatography. To the protein solution prepared from <0.5 or >0.5 g wet cells, 1 or 2 mL of the cOmplete His-Tag purification resin (Roche) equilibrated with the solubilization buffer 2 was added, respectively, and the mixture was rotated at room temperature for more than 1 h. The resin was placed in a column and washed with solubilization buffer 2 until the absorbance at 280 nm (A_280_) of the eluate was <0.1 (typically 10 cv). The protein was eluted with solubilization buffer 2 supplemented with 200 mM imidazole (15–30 cv).

The protein concentration of solubilized ECD or purified ECD-His was estimated by measuring the A_280_. A molar extinction coefficient of 36,370 was used for the ECD variants, except for the W24A and W28A variants, whose molar extinction coefficient was 30,680 (27). The protein solution was diluted by the solubilization buffer 1 or 2 to 0.5 mg/mL for ECD or 0.1 mg/mL for ECD-His, and the solution was dialyzed against the dialysis buffer 1 containing 20 mM Tris-HCl (pH 8.2) and 50 mM NaCl with 20 times as much volume as the inner solution at 4 °C overnight. The outer buffer was exchanged for dialysis buffer 2, which contained 20 mM Tris-HCl (pH 8.2) and 60 mM NaCl, and the dialysis was continued at 4 °C for a day.

After the dialysis, the inner solution was centrifuged at 27,143 ×g and 4 °C for 30 min. The supernatant was collected. The refolded proteins were purified using anion-exchange chromatography. To the dialysis inner solution, 1 mL of the Q Sepharose Fast Flow Resin (Cytiva) equilibrated with the QA buffer containing 25 mM Tris-HCl (pH 8.0) was added, and the mixture was rotated at 4 °C for 20 min. The resin was placed in a column and washed with QA buffer until the A_280_ of the eluate was <0.1 (typically 10 cv). The protein was then eluted sequentially with QA buffer supplemented with 100, 200, and 300 mM NaCl to obtain eluates 1 (4 cv), 2 (20–30 cv), and 3 (5–48 cv). The eluates 1-3 were mixed and concentrated with Amicon Ultra-15 (Merck Millipore, MWCO 3 K) to protein concentrations of 44–162 µM.

In the case of the D46V variant, the protein did not bind to the Q Sepharose resin, and the pass-through was purified by Ni^2+^ affinity chromatography. To the protein solution prepared from 0.75 g wet cells, 2 ml of the cOmplete His-Tag purification resin (Roche) equilibrated with the Tris buffer 4 containing 25 mM Tris-HCl (pH 8.0) and 100 mM NaCl was added. The mixture was stirred at 4 °C for 20 min. The resin was placed in a column, washed with Tris buffer 4 (5 cv), and washed with a buffer containing 20 mM Tris-HCl (pH 7.4), 100 mM NaCl, and 30 mM imidazole (5 cv). The protein was then eluted by an elution buffer containing 20 mM Tris-HCl (pH 7.4), 100 mM NaCl, and 200 mM imidazole (13.5 cv). The eluate was concentrated with Amicon Ultra-15 (Merck Millipore, MWCO 3 K) to a protein concentration of 60 µM.

Final purification was performed using size-exclusion chromatography (SEC). The concentrated protein solution was loaded into a HiLoad 16/600 Superdex 200 pg column (Cytiva) connected to an NGC Quest 10 Plus system (Bio-rad) placed in a 4 °C chamber. A phosphate buffer containing 10 mM NaPi (pH 6.4) and 50 mM NaCl and the standard buffer containing 20 mM Tris-HCl (pH 7.4) and 100 mM NaCl were used as running buffers for the purification of ECD and ECD-His, respectively. The flow rate was 0.15 mL/min, injection volumes were 0.7–5.6 mL, and the fractional volume was 5 mL.

The yields of the purified ECDs were 12 mg and 30 mg per liter of culture for the 8-mer and monomer, respectively. The yields of purified ECD-His were 6–8 mg and 8–11 mg per liter of culture for the 8-mer and monomer, respectively. The yields of the purified monomeric variants of ECD-His (W24A, W28A, and N87H) were 25–41 mg per liter of culture. The yields of the other ECD-His variants varied from 1–35 mg and 1–13 mg per liter of culture for the 8-mer and monomer, respectively.

### SEC-MALS analysis of ECD-His

The SEC-MALS experiment was carried out at room temperature using an Alliance HPLC system (Waters) combined with a DAWN HELEOS II detector (Wyatt) and a Superdex 200 Increase 10/300 GL column (Cytiva) equilibrated with the standard buffer. An ECD-His solution with a concentration of 52 µM was analyzed in the standard buffer. The flow rate was 0.5 mL/min. The injection volume was 0.2 mL. The data were processed using ASTRA 6.1 software (Wyatt) with a dn/dc value of 0.185 ml/g, which is typical for non-glycosylated proteins (28).

### Crystallization and structure determination of ECD

The second SEC eluate of ECD, with a peak at 102 mL, was collected and dialyzed against a MES buffer containing 10 mM MES (pH 6.4) and 50 mM NaCl. The protein was concentrated with an Amicon Ultra-4 (Merck Millipore, MWCO 3 K) to a concentration of 726 µM. Initial crystallization screening was performed by the sitting-drop vapor-diffusion method using commercially available screening kits such as Crystal Screening I & II (Hampton Research) and Wizard I & II (Rigaku) with a Protein Crystallization System 2 (PXS2) at the SBRC in KEK (29). Droplets containing equal volumes of reservoir solution (0.5 µL) and protein solution (0.5 µL) were incubated with 200 µL of each reservoir solution at 20 °C. The best crystal was obtained in a solution containing 100 mM Tris-HCl (pH 8.5) and 300 mM magnesium formate within one week. The crystal was soaked in an increased concentration of magnesium formate with 30% v/v glycerol and frozen in liquid nitrogen. Diffraction data were collected at 100 K at a wavelength of 1.0 Å on the SPring-8 BL32XU using the Eiger X 9M detector. The datasets (360° in 0.1° steps) were processed and scaled using the XDS program package (30) and Aimless in CCP4 suite (31). The crystal belongs to the space group *I422*, with one molecule in the asymmetric unit. Phase information was obtained by molecular replacement with the PHASER MR (Phaser crystallographic software) (32) using the crystal structure of rMPZ-ECD (PDB ID 1NEU) as the search model. The replacement model was manually constructed using COOT (33) and refined to a resolution of 2.1 Å using the PHENIX package (34). Extra electron density was found on the surface of ECD adjacent to the *trans* interaction site. This might correspond to a glycerol molecule used as the cryoprotecting agent. The final model contained the ECD residues I1–V121 and a glycerol. Data collection and refinement statistics are presented in Table S2. The figures were prepared using PyMOL (Schrödinger).

### Preparation of ECD-attached NDs

The plasmid for the expression of MSP1D1 was obtained from Addgene, and the MSP1D1 protein was produced and purified as previously described (11). Powders of 1,2-dimyristoyl-*sn*-glycero-3-phosphocholine (DMPC) and 1,2-dioleoyl-*sn*-glycero-3-[(N-(5-amino-1-carboxypentyl)iminodiacetic acid)succinyl]-Ni^2+^ (DOGS-NTA-Ni^2+^) (Avanti Polar Lipids) were dissolved in chloroform and mixed in a glass tube at a molar ratio of 62:8. Chloroform was evaporated by blowing a gentle stream of nitrogen gas while rotating the glass tube to prepare lipid films on the glass surface. Lipid films were dried overnight using a lyophilizer (SP VirTis). A solution containing 100 mM sodium cholate and 100 mM NaCl was added to the lipid films to achieve the final total lipid concentration of 70 mM (the concentrations of DMPC and DOGS-NTA-Ni^2+^ were 62 and 8 mM, respectively), and lipids were dissolved by immersing the glass tube in a water bath at 50–70 °C and vortexing five times repeatedly. For the preparation of nanodiscs (NDs), an 8 mM lipid mixture, 114 µM MSP1D1, and a total of 15 mM sodium cholate were prepared in the standard buffer and rotated at room temperature for 2 h. The solution was dialyzed at room temperature against the standard buffer with 417 times as much volume as the inner solution. After more than 6 h, the outer buffer was replaced with fresh buffer, and dialysis was continued for more than 6 h. The prepared NDs were purified by SEC using a Superdex 200 Increase 10/300 GL column (Cytiva) connected to an NGC Quest 10 Plus system (Bio-rad) and the standard buffer as the running buffer at 4 °C. The flow rate was 0.25 mL/min. The injection volume was 0.25 ml, and the fractional volume was 1 mL. The concentration of the purified ND was determined using A_280_ and a molar extinction coefficient of 36,400, originating from two MSP1D1 molecules per ND (35).

To prepare ECD-His-attached NDs, we used the 8-meric ECD-His included in the first elution peak of the SEC purification, except for the W24A, W28A, and N87H variants, which existed only as monomers. NDs and ECD-His were mixed in the standard buffer at a molar ratio of 1:8 (four ECD-His molecules per side of an ND). The final concentrations of NDs and ECD-His ranged from 0.2–1.4 µM and 1.4–11 µM, respectively. The mixture was incubated in a heat block at 30 °C for 20 min. and centrifuged at 17,800 ×g and 25 °C for 3 min. The supernatant was concentrated with Amicon Ultra-4 (Merck Millipore, MWCO 3K) to approximately 250 µL and analyzed by SEC using a Superdex 200 Increase 10/300 GL column (Cytiva) and the standard buffer at 4 °C. The flow rate was 0.25 mL/min. The fractional volume was 1 mL. The eluates were then used for electron microscopy.

In an inhibition experiment to analyze the contribution of the inter-ECD interactions in membrane-stacking, a sample containing NDs, ECD-His, and the 8-meric ECD (without a histidine-tag) with a molar ratio of 1:8:80 was prepared and analyzed by SEC as described above. To confirm the contribution of the interaction between Ni^2+^-chelated lipids and ECD-His, a mixture of ND, ECD-His, and 250 mM imidazole was prepared and analyzed by SEC in the standard buffer supplemented with 250 mM imidazole.

### Electron microscopic studies of ECD-attached NDs (Nanomyelin)

The ND-ECD-His variant complexes eluted at the void volume (8.2 mL) were subjected to electron microscopy (EM) studies. For the NDs linked to the W24A, W28A, and N87H variants, SEC eluates in the peak at 10–11 mL were subjected to EM studies. The concentration of the ND-ECD-His variant complex was determined using A_280_ and molar extinction coefficients of 327,360 or 281,840 (for the W24A and W28A variants), which originated from a unit of nanomyelin composed of two MSP1D1 and eight ECD-His molecules. The concentration of the ND-ECD-His variant complex was adjusted to 0.04 μM using the standard buffer. The samples were adsorbed onto thin carbon films supported by copper mesh grids (Nisshin-EM), whose surfaces were hydrophilized by glow discharge under low pressure. The samples were negatively stained with an EM stainer (Nisshin EM), blotted, and air-dried. The samples were observed under a JEM-1230 transmission electron microscope (JEOL) at an acceleration voltage of 120 kV. Images were recorded using a F114T CCD camera (TVIPS).

### CD analyses of ECD-His

CD experiments were performed using a J-720 spectrometer (JASCO) equipped with a Peltier-type thermal controller. The concentrations of the ECD-His variants were adjusted to 6–22 µM in the standard buffer. The temperature-dependent change in the CD signal at 217 nm was monitored for each ECD-His variant in a quartz cuvette with a 0.2 cm path length. The temperature was increased from 20 to 90 °C at intervals of 1 °C and a gradient of 3 °C per min. Midpoint temperatures of denaturation (T_m_s) were determined using CDpal software (13).

### Data and code availability

Atomic coordinates for human MPZ-ECD have been deposited in the PDB (https://www.rcsb.org/) with an ID of 8IIA.

## Supporting information

Supplemental results, figures, and tables

## Acknowledgements and funding sources

The authors thank M. Henmi and M. Ito for technical assistance. We also thank the beamline staff at SPring-8 BL32XU, Photon Factory BL-1A, BL-17A and Swiss Light Source PXI(X06SA). This work was supported by JSPS KAKENHI Grants 18K06601 and 21K06515 (to M.S.), JST CREST Grant Number JP18071859 (to K.M.), and the RIKEN pioneering project “Dynamic Structural Biology” (to H.T.). This crystallographic study was supported by Platform Project for Supporting Drug Discovery and Life Science Research and Research Support Project for Life Science and Drug Discovery (Basis for Supporting Innovative Drug Discovery and Life Science Research (BINDS)) from AMED under Grant Numbers JP21am0101070 and JP22ama1210001j001 (support numbers #2101 and #4101). Diffraction experiments at SLS were covered under project ID 20191094 and 20191134.

## Author contributions

M.S. planned the experiments and M.S., M.T., and K.M. wrote the manuscript. M.S., M.M., and H.T. prepared the ECD samples. M.S. and M.T. performed SEC-MALS experiments. M.T. performed crystallographic analysis. M.S. prepared the NDs and nanomyelin. M.S. and K.M. performed the EM experiments.

