## Supplemental results, figures, and tables for "Structural bases for the Charcot–Marie–Tooth disease induced by single amino acid substitutions of myelin protein zero"

### Supplemental Information

### Supplemental Results

#### Structural differences between the multimeric hMPZ-ECD and rMPZ-ECD

Out of the amino acid substitution sites from the human to rat ECDs (H10Y, R16Q, and R77S), R/Q16 and R/S77 were located at the *cis* and head-to-head interfaces, respectively (Fig. 2B, 2D, and S3A). R16 of human ECD formed a cation-π interaction and a hydrogen bond with Y4 and S22 of the neighboring ECD, respectively. In contrast, Q16 in the rat ECD formed a water-mediated hydrogen bond with S22 in the facing ECD (Fig. S3B). R77 of the human ECD makes van der Waals contact with the R77 of the opposing ECD. The electric repulsion between the guanidinium groups of R77 may be attenuated by the cation-π interaction between R77 and W78 (Fig. S3C). In contrast, S77 of the rat ECD interacted with the facing S77 via three water-mediated hydrogen bonds (Fig. S3C). The quantitative effects of these substitutions on each interaction remain unclear.

**Supplemental Figures and Tables**

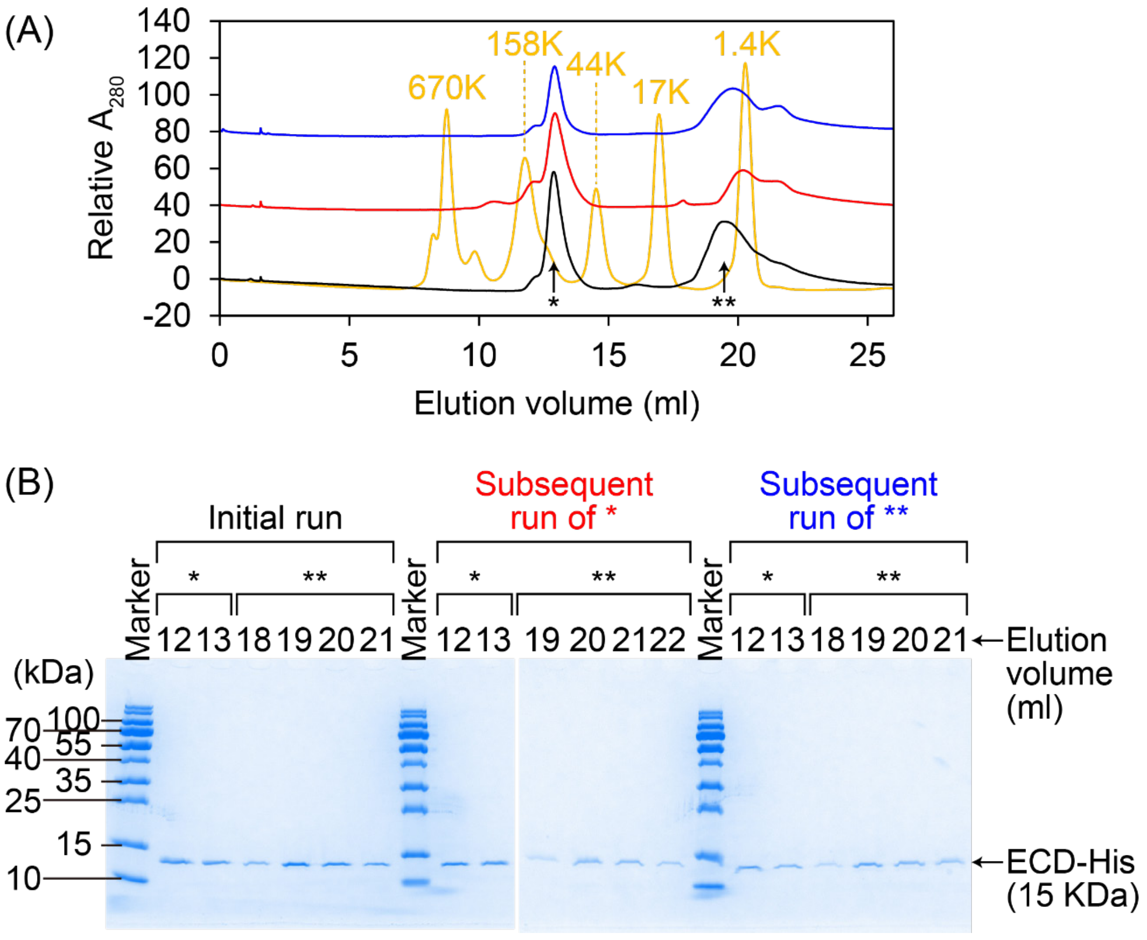


**Figure S1.** SEC analyses of ECD-His. (A) Profiles of the initial SEC experiment for ECD-His after the purification by the anion exchange chromatography (black), a subsequent SEC experiment for ECD-His contained in the first elution peak of the initial SEC experiment shown by the arrow with * (red), and a subsequent SEC experiment for ECD-His contained in the second elution peak of the first SEC experiment shown by the arrow with ** (blue). Both the red and blue runs produced two major elution peaks with equivalent elution volume as * and ** in the initial run. The orange line shows the profile of the MW standards. The concentration of the injected ECD-His was 50 μM (black), 37 μM (red), and 47 μM (blue). The experiments were performed at 4 °C in the standard buffer using a Superdex 200 Increase 10/300 column (Cytiva). (B) Results of the SDS-PAGE analyses of the SEC eluates. Both the first and second peaks solely contained ECD-His.

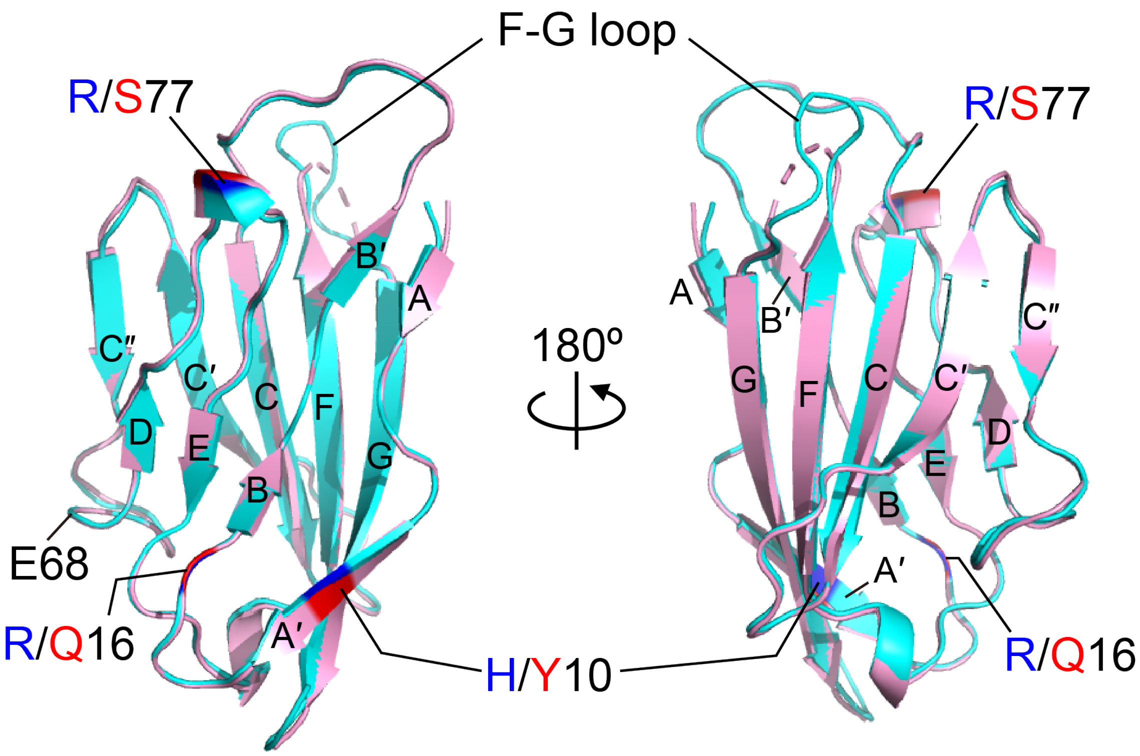


**Figure S2.** Superimposed crystal structures of human MPZ-ECD (cyan) and rat MPZ-ECD (pink: PDB ID 1NEU). ECD forms a typical immunoglobulin-fold, and β-strands are labeled after the immunoglobulin variable domain. Three substitution sites are shown in blue (human) and red (rat). The CMT-related E68V substitution site is also labeled. The F-G loop that was disordered in the rat structure was observed in the human structure. The RMSD of the two structures is 0.387 Å.

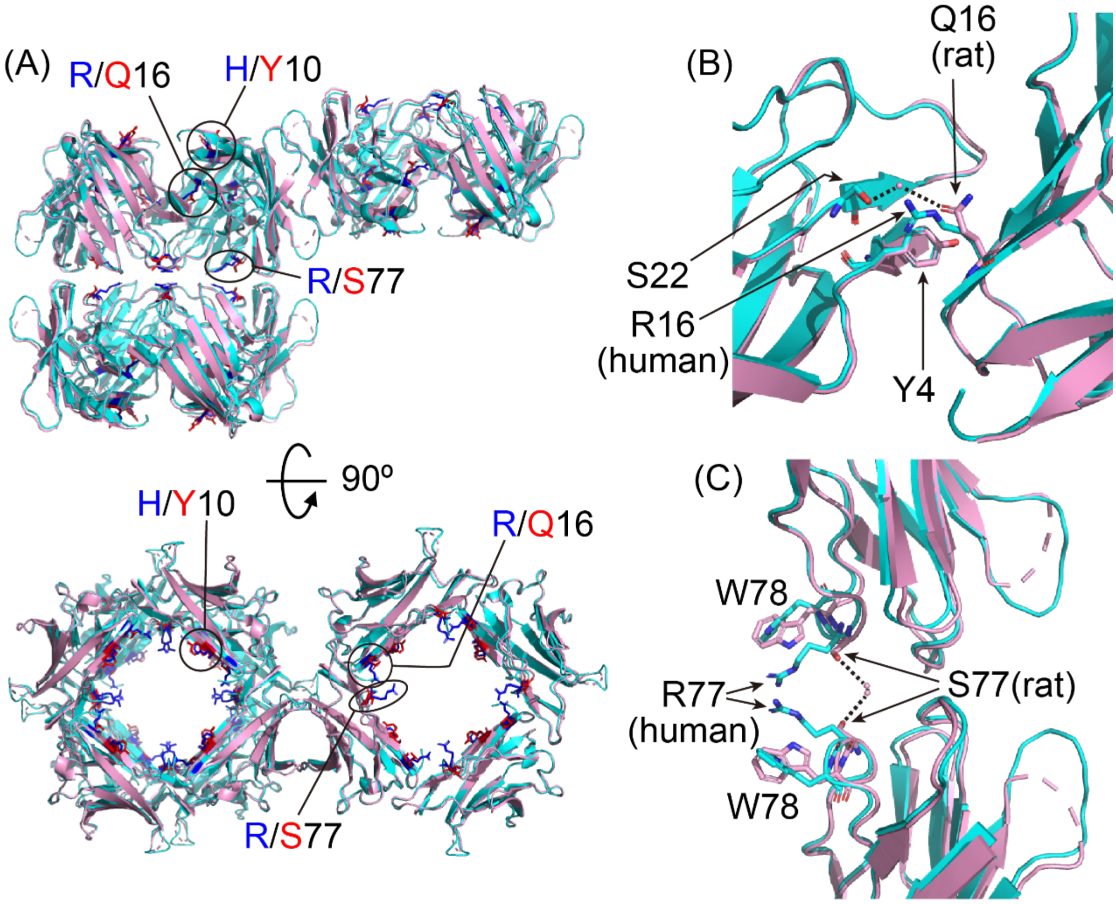


**Figure S3.** Superimposed crystal structures of the multimeric forms of human MPZ-ECDs (cyan) and rat MPZ-ECDs (pink: PDB ID 1NEU). (A) Overview of the multimerized structures of MPZ-ECDs. Three substitution sites are shown by sticks and colored blue and red for human and rat proteins, respectively. (B) Expanded view of the *cis* interaction site between ECDs. R16 in human ECD forms interactions with Y4 and S22 of the facing ECD, while Q16 interacts with S22 via water-mediated hydrogen-bonds (dotted lines). (C) Expanded view of the head-to-head interaction site between ECDs. R77 in human ECD directly interacts with R77 from the facing ECD, while the S77 residues in rat ECDs are connected with each other via water-mediated hydrogen-bonds (dotted lines).

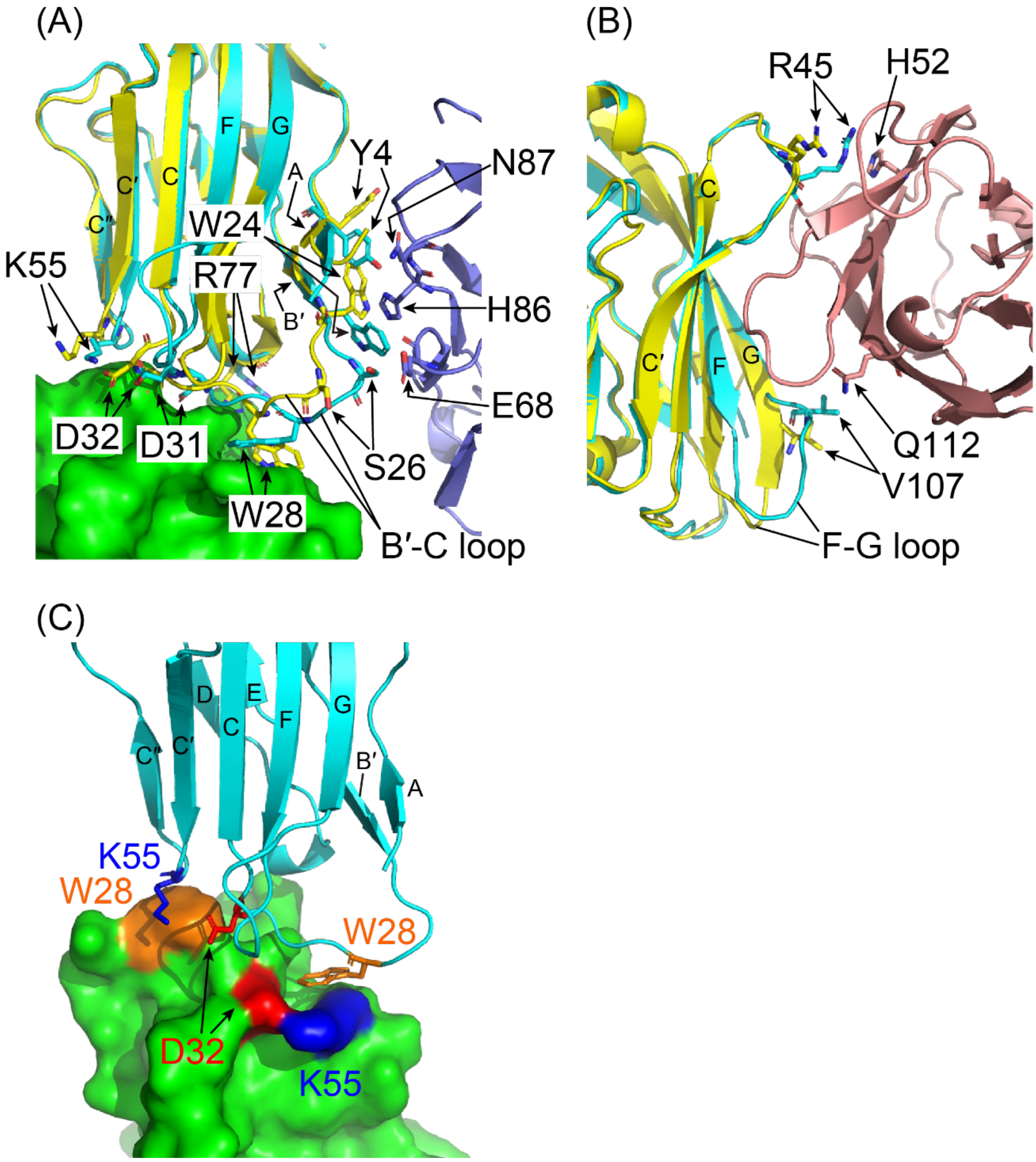


**Figure S4.** Superimposed crystal structures of the multimerized human MPZ-ECDs (cyan, slate blue, green, and salmon pink) and the monomeric human MPZ-ECD (yellow: PDB ID 3OAI). (A) Expanded view of the *cis* and head-to-head interaction sites between ECDs. The B′-C loop form both the *cis* and head-to-head interaction sites in the multimerized structure (cyan, slate blue, and green), while these residues in the monomeric structure are not suitably arranged for the inter-ECD associations (yellow). (B) Expanded view of the *trans* interaction site between ECDs. V107 on the F-G loop and R45 interact with Q112 and H52 from the facing ECD in the multimerized structure (cyan and salmon pink), respectively. On the other hand, these residues of the monomeric ECD (yellow) are apart from the adjacent ECD when the monomeric ECD is superimposed on the multimeric ECDs. (C) Expanded view of the head-to-head interaction site between ECDs. D32 and K55 make an electrostatic interaction with each other to form a cavity to accommodate W28 protruded from the interacting ECD.

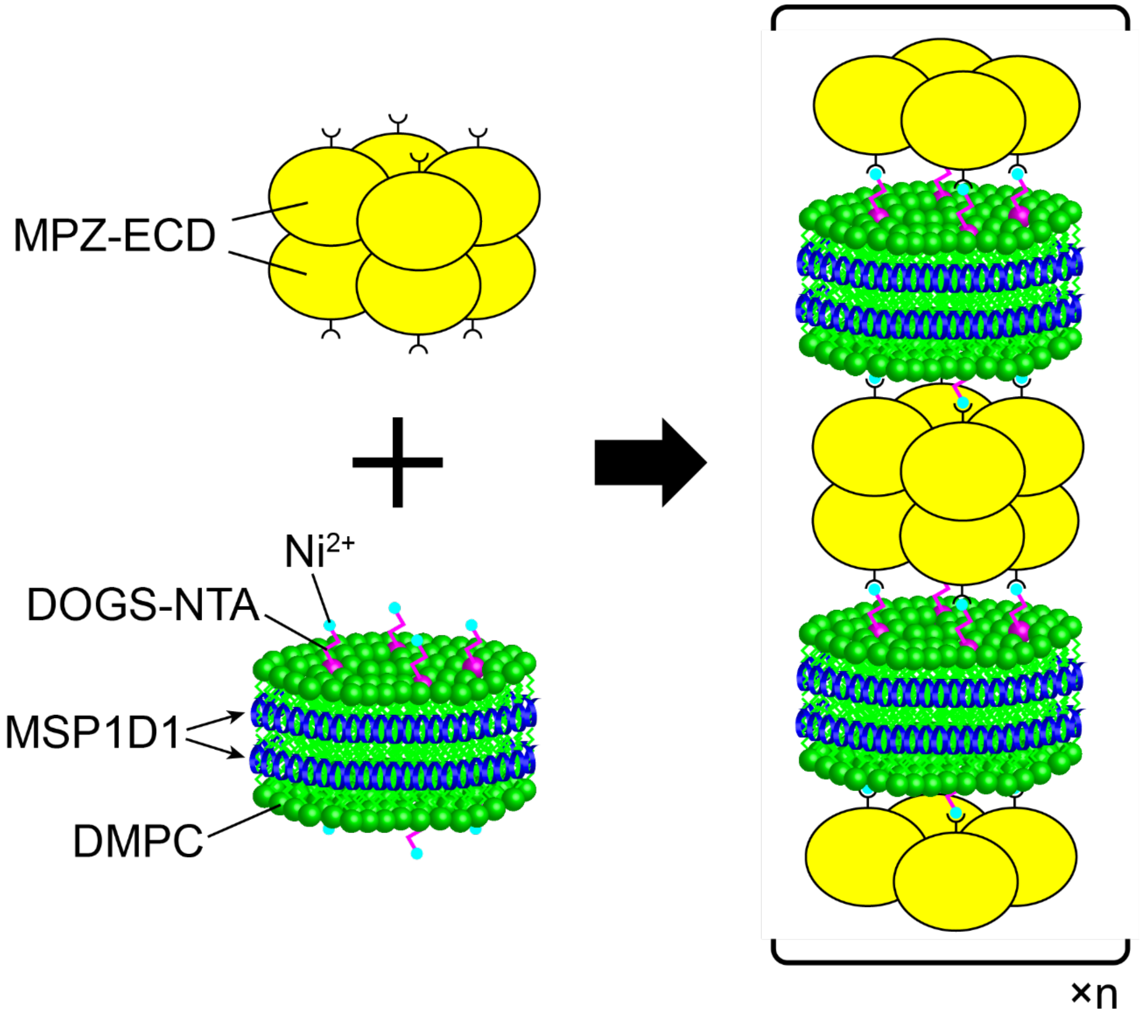


**Figure S5.** Schematic illustration of NDs stacked by the ECD-His-8-mer (nanomyelin). Nanomyelin were prepared by mixing the ECD-His-8-mer and NDs. In each ND, hydrophobic edge of the lipid bilayer composed of DMPC (green) and DOGS-NTA (magenta) is surrounded by two MSP1D1 molecules (blue). ECD-His-8-mers (yellow) are attached to each side of an ND via the interaction between a histidine-tag of ECD-His and a $Ni^{2+}$ ion (cyan) chelated by DOGS-NTA and glue NDs.

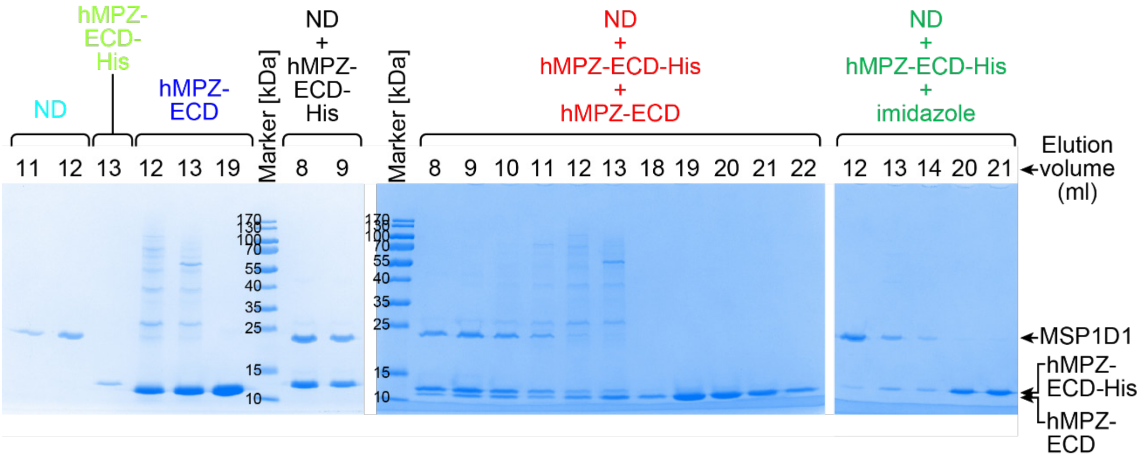


**Figure S6.** SDS-PAGE analyses of nanomyelin. Proteins contained in each elution peak of the SEC (shown in Fig. 3) were analyzed by SDS-PAGE. The colors used in this figure correspond to the SEC charts with the same colors in Fig. 3.

**Table S1.** Amino acid sequences of ECD and ECD-His

| Construct | Amino acid sequence |
|---|---|
| ECD | MGIVVYTDREVHGAVGSRVTLHCSFWSSEWVSDDISFTWRYQPEGGRD AISIFHYAKGQPYIDEVGTFKERIQWVGDPRWKDGSIVIHNLDYSDNGTFT CDVKNPPDIVGKTSQVTLYVFEKV |
| ECD-His | MGIVVYTDREVHGAVGSRVTLHCSFWSSEWVSDDISFTWRYQPEGGRD AISIFHYAKGQPYIDEVGTFKERIQWVGDPRWKDGSIVIHNLDYSDNGTFT CDVKNPPDIVGKTSQVTLYVFEKVPTRHHHHHH |

The N-terminal artificially inserted residues for expression and the linker residues connecting the C-terminus of ECD and the histidine-tag are colored in red. The histidine-tag is shown in green.

**Table S2.** Statistics from the crystallographic analysis of ECD

| | |
|---|---|
| **Data collection** | |
| Synchrotron Beamline | SPring-8 BL32XU |
| Wavelength (Å) | 1.00 |
| Space group | *I422* |
| Cell dimensions<br>*a*, *b*, *c* (Å) | 88.5, 88.5, 88.9,<br>90 = 90 = 90 |
| Resolution (Å) | 44.29–2.09 (2.14–2.09) |
| *R*pim | 0.014 (0.634) |
| $I/\sigma(I)$ | 13.9 (1.1) |
| Completeness (%) | 100 (99.5) |
| $CC_{1/2}$ | 99.8 (85.1) |
| Multiplicity | 26.8 (27.4) |
| Total observations | 291217 |
| Unique observations | 10861 (815) |
| **Refinement** | |
| Resolution (Å) | 2.09 |
| Number of reflections | 10343 |
| $R_{work}/R_{free}$ (%) | 21.9/23.3 |
| Number of atoms | 1040 |
| Protein | 992 |
| Solvent | 48 |
| Average B-factors (Å$^2$) | 53.2 |
| Protein | 53.1 |
| Solvent | 55.9 |
| **RMSDs** | |
| Bond lengths (Å) | 0.005 |
| Bond angles (°) | 0.700 |
| **Ramachandran** | |
| Ramachandran favored (%) | 96.6 |
| Ramachandran outliers (%) | 0 |
| **PDB ID** | 8IIA |

*Values in parentheses correspond to highest-resolution shells.
